# PROTECTIVE ROLE FOR SMOOTH MUSCLE CELL HEPCIDIN IN ABDOMINAL AORTIC ANEURYSM

**DOI:** 10.1101/2021.07.30.454447

**Authors:** P Loick, G Mohammad, I Cassimjee, A Chandrashekar, P Lapolla, A Carrington, A Handa, R Lee, S Lakhal-Littleton

## Abstract

**Rationale:** Hepcidin (HAMP) is a hormone produced primarily in the liver. It controls systemic iron homeostasis by inhibiting the iron exporter ferroportin (FPN) in the gut and spleen, respective sites of iron absorption and recycling. HAMP and FPN are also found ectopically in tissues not involved in systemic iron homeostasis. The physiological functions of ectopic HAMP and FPN are only just beginning to be uncovered. We observed that HAMP expression is markedly increased in smooth muscle cells (SMCs) of abdominal aortic aneurysms (AAA), both in patients and in an experimental mouse model of AAA.

**Objective:** To understand the role of SMC-derived HAMP in the pathophysiology of AAA.

**Methods and Results:** We generated mice harbouring an inducible, SMC-specific deletion of the *hamp* gene. We then applied the experimental model of AAA and simultaneously induced deletion of *hamp* in SMCs. We found that these mice developed large aneurysms and had greater incidences of rupture and of fatal dissection than mice with intact *hamp* in SMCs. A similar phenotype was observed in mice harbouring an inducible SMC-specific knock-in of HAMP-resistant FPNC326Y. Additionally, we observed that expression of Lipocalin-2 (LCN2), a protein known to promote AAA, was suppressed in AAA tissue from patients and from mice with intact *hamp* in SMCs, but not in mice lacking *hamp* in SMCs. Treatment of these mice with a LCN2-neutralising antibody protected them from the otherwise detrimental effects of loss of *hamp* in SMCs.

**Conclusions:** The present study demonstrates that the rise in SMC-derived HAMP within the aneurysm tissue is protective in the setting of AAA, and that such protection involves the cell-autonomous action of HAMP, and suppression of local LCN2. These findings are the first example of a protective role for ectopic HAMP in disease. They expand understanding of the multifaceted functions of HAMP outside the liver.

## INTRODUCTION

An abdominal aortic aneurysm (AAA) is an abnormal dilatation of the abdominal aorta, defined as an aortic dimeter of >3cm. Left untreated, an AAA gradually expands and can potentially lead to a catastrophic rupture. Approximately ∼200,000 people globally demise yearly from an AAA^1^. Patients with AAA are offered an elective repair at 5.5cm to prevent a future rupture event. In many countries screening programs exist for at-risk populations^2^. Traditional risk factors for developing an AAA include cigarette smoking, male gender, age > 65 and a family history of AAA. In addition to the risk of AAA rupture, these patients also are at increased risk of mortality from cardiovascular complications^3^.

Hepcidin or human antimicrobial peptide (HAMP) is the master hormone of systemic iron homeostasis. It is primarily produced in the liver and controls systemic iron homeostasis by endocrine inhibition of the iron exporter ferroportin (FPN) in duodenal enterocytes and splenic reticuloendothelial macrophages, respective sites of iron absorption and recycling^4,5^. HAMP and FPN are also found ectopically in tissues with no recognized role in systemic iron homeostasis, including the heart, the brain, kidney, and pulmonary arterial smooth muscle cells^6-10^. In the heart and pulmonary arterial smooth muscle cells, work from this laboratory has shown that HAMP acts cell-autonomously to regulate intracellular iron levels, and that such regulation is important for the normal physiological function of the respective tissue^6,7,10^. HAMP is also induced by inflammation, accounting for iron deficiency of chronic diseases^11^. Raised plasma levels of HAMP have been reported in AAA patients^12^. Additionally, there is an association between iron deficiency anaemia and increased AAA size, and between excess iron deposition within the aneurysm tissue and markers of local oxidative damage^13,14^. However, the contribution of ectopic HAMP tissue expression to raised plasma HAMP, and its precise role in the pathophysiology of AAA remain unknown. In this study, we report that both in patients and in an experimental mouse model of AAA, HAMP expression is markedly raised in SMCs within the aneurysm tissue. To probe the disease-significance of raised HAMP in SMCs without the confounding effects of altered systemic iron homeostasis, we generated novel mice harbouring an inducible SMC-specific knockout of the *hamp* gene or an inducible SMC-specific knock-in of a HAMP-resistant FPN isoform FPNC326Y.

## METHODS

### Participants, blood sampling, and AAA growth rate measurement

Details regarding the Oxford Abdominal Aortic Aneurysm (OxAAA) study cohort and recruitment process have been published^15^. In brief, this single centre prospective study (Ethics Ref: 13/SC/0250) recruited patients in the National Health Service (UK). Each participant gave written informed consent. Baseline assessments were performed and included demographic and risk factor data, antero-posterior AAA diameter measurements by ultrasound imaging, and fasting venous blood sampling. Platelet-poor plasma (PPP) was prepared immediately after blood collection at room temperature using two-staged centrifugation (1^st^ stage: 1300gx12min; 2^nd^ stage: 2500gx15min) as previously described^15^ These were stored at -80°C for subsequent analysis. Prospective AAA annual growth were calculated based on the antero-posterior diameter (APD) measurements in the subsequent yearly AAA monitoring ultrasound scan: (ΔAPD/APD at baseline)/(number-of-days-lapsed/365days)^16^.

### Intra-operative tissue biopsy of omental artery and aneurysm tissue

Intraoperative biopsies of paired tissue samples were performed as previously described^17^. Tissue samples were prepared immediately in the operating theatre. During AAA surgery, a wedge of abdominal omentum containing a segment of omental artery was identified and biopsied en-bloc. The omental artery segment was cleared of peri-vascular tissue and snap-frozen. A longitudinal strip of the aneurysm wall along the incision was then excised. The aneurysm tissue was stripped off the peri-vascular tissue and mural thrombus. The tissue at the maximal dilatation was isolated, divided into smaller segments, and snap-frozen for subsequent analysis. Omental artery and aneurysm wall specimens were also suspended in paraformaldehyde (4%), and later embedded in paraffin moulds. These were sectioned for use in histology and immunohistochemistry experiments. Non-aneurysmal aortas were procured from deceased transplant donors. These were sourced from the Oxford Transplant Biobank (Ethics Ref: 14/SC/1211), which curates tissue samples collected from the organ donors. As with aneurysmal and omental tissue, fragments were snap-frozen and embedded in paraffin for sectioning.

### Mice

All animal procedures were compliant with the UK Home Office Animals (Scientific Procedures) Act 1986 and approved by the University of Oxford Medical Sciences Division Ethical Review Committee.

The conditional hepcidin *hamp* and ferroportin *fpnC326Y* floxed alleles (fl) were generated as described previously^6,7,10^. Targeting of these conditional alleles to SMCs was achieved by crossing to mice transgenic for Y chromosome-linked tamoxifen-inducible CreER^T2+^ gene under the control of the smooth muscle myosin heavy chain polypeptide 11 (myh11) promoter (SMMHC-CreER^T2+^)^18^. Male hamp^fl/fl^, SMMHC-CreER^T2+^ and fpnC326Y^fl/fl^, SMMHC-CreER^T2+^ animals were treated with an intraperitoneal injection of tamoxifen, for timed deletion of hamp, and knock-in of fpnC326Y respectively. Male hamp^fl/fl^ and fpnC326Y^fl/fl^ animals were used as respective littermate controls, and also treated with tamoxifen at the same timepoint. To induce AAA in mice, recombinant Angiotensin II (AngII) was delivered at a dose of 1 μg/kg/min over 7 days, using subcutaneously-implanted slow release Alzet osmotic minipumps (Alzet 1004) as described previously^19,20^. Animals were culled at the 1 week timepoint for collection of plasma and tissues. Animals that died before the 1 week timepoint were subject to post-mortem examination to ascertain the cause of death. Lipocalin (LCN2)-neutralising antibody MAB1857 (R&D systems) was infused intravenously at 4mg/Kg.

### Quantitative PCR

Total RNA extraction and cDNA synthesis were carried out as previously described^6,7,10^. Gene expression was measured using Applied Biosystems Taqman gene expression assay probes for Hamp, Transferrin receptor TfR1, Divalent metal transporter Dmt1, Lipocalin-2 Lcn2 and house-keeping gene β-Actin (Life Technologies, Carlsbad, CA). The CT value for the gene of interest was first normalised by deducting CT value for β-Actin to obtain a delta CT value. Delta CT values of test samples were further normalised to the average of the delta CT values for control samples to obtain delta delta CT values. Relative gene expression levels were then calculated as 2-^delta deltaCT.^

### Histology and Immunohistochemistry

Formalin-fixed paraffin-embedded tissues were sectioned into 0.5-µm sections and stored at room temperature. Sections were deparaffinized using xylene and then were rehydrated in ethanol. For imaging of aortas, slides were stained for Elastin using Elatin Stain kit (HT25A, Sigma-Aldrich) as per manufacturer’s instructions. For DAB-enhanced Perls iron stain, slides were immersed for 1 h in 1% potassium ferricyanide in 0.1 M HCl buffer and then were stained with DAB chromogen substrate. All slides were counterstained with hematoxylin and then were visualized using a standard brightfield microscope. For immunostaining, slides were subjected to heat-activated antigen retrieval in citrate buffer (pH 6), then stained with rabbit polyclonal anti-HAMP antibody (ab30760, Abcam) at 1/40 dilution. For non-fluorescent immunostaining, EnVision+ System-HRP (K4011, Dako) was used as secondary antibody as supplied. For fluorescence immunostaining, Alexa 488-conjugated anti-rabbit antibody (ab150073, Abcam) was used as secondary antibody at a dilution of 1/500, and slides were also counter-stained with Cy3-conjugated anti-mouse alpha smooth muscle Actin SMA-α antibody (A2547 Sigma-Aldrich) at 1/500. Following counter stain with DAPI, coverslips were visualised using a Fluoview FV1000 confocal microscope (Olympus).

### Iron quantitation

Plasma iron levels were determined using the ABX-Pentra system (Horiba Medical, CA). For quantitation of iron in tissues, animals were first perfused with saline to remove blood from tissues. Determination of total elemental iron in tissues was carried out by inductively coupled plasma mass spectrometry (ICP-MS) as described previously^6,7,10^.

### Enzyme-Linked Immunoassay

Human HAMP was measured in plasma by ELISA DHP250 (R&D Systems). Mouse HAMP was measured in plasma by ELISA LS-F11620 (Ls-Bio).

### Statistics

Values are shown as mean ± Standard error of the mean (SEM). Paired comparisons were performed using Student’s t test. Multiple comparisons were drawn using ANOVA. Post hoc tests used Bonferroni correction.

## RESULTS

### HAMP is elevated in SMCs of AAA in patients and in mice

We examined the expression of HAMP in aneurysm tissue excised from AAA patients. We found that HAMP protein was markedly abundant in the aneurysm tissue, in contrast to abdominal aortic tissue from non-AAA healthy controls (HC), and to control vascular tissue (omental artery) from AAA patients (Figure 1A). Co-localisation staining with SMC marker; smooth muscle cell actin alpha (SMA-α); demonstrated that most HAMP protein in the aneurysm tissue localised to SMCs (Figure 1A). Relative differences in HAMP protein abundance were replicated at the mRNA level, confirming that HAMP protein present in the aneurysm tissue is the product of local gene expression (Figure 1B). When we examined plasma HAMP levels, we found that these were significantly higher in AAA patients than in HCs (Figure 1C). To explore these findings further, we used the Ang-II-infusion experimental mouse model, previously shown to induce AAA in mice even against a wild type background^19,20^. We found that this model resulted in AAA in a small proportion of wild type mice after 1 week of Ang-II delivery. Where AAA was present, we detected a marked upregulation of HAMP in SMCs with the aneurysm tissue (Figure 1D). To further assess the contribution of SMCs to raised HAMP levels, we applied the Ang-II-infusion model to mice harbouring a conditional tamoxifen-inducible deletion of the *hamp* gene specifically in SMCs (Hamp ^fl/fl^, SMMHC-CreER^T2+^). In control mice (Hamp ^fl/fl^), Ang-II treatment resulted in ∼31 fold increase in hamp mRNA in the abdominal aorta (Figure 1E). However, this increase was only 5 fold in Hamp^fl/fl^, SMMHC-CreER^T2+^ mice, demonstrating that SMCs are the dominant source of locally-produced HAMP in the aneurysm tissue (Figure 1E). Additionally, plasma HAMP levels were raised by Ang-II treatment in Hamp^fl/fl^ mice but not in Hamp ^fl/fl^, SMMHC-CreER^T2+^ mice, demonstrating that the rise in SMC-derived HAMP in the aneurysm tissue is sufficient to spill into the circulation (Figure 1F). Of note, *hamp* gene expression was not raised by Ang-II in the liver in either sets of mice, supporting the notion that raised plasma HAMP in the setting of AAA reflects its increased release from SMCs of the aneurysm tissue rather than from hepatocytes (Figure 1G). Together, these results demonstrate that HAMP is elevated in the SMCs within AAA and that this rise contributes to raising plasma HAMP levels.

**Figure 1.**
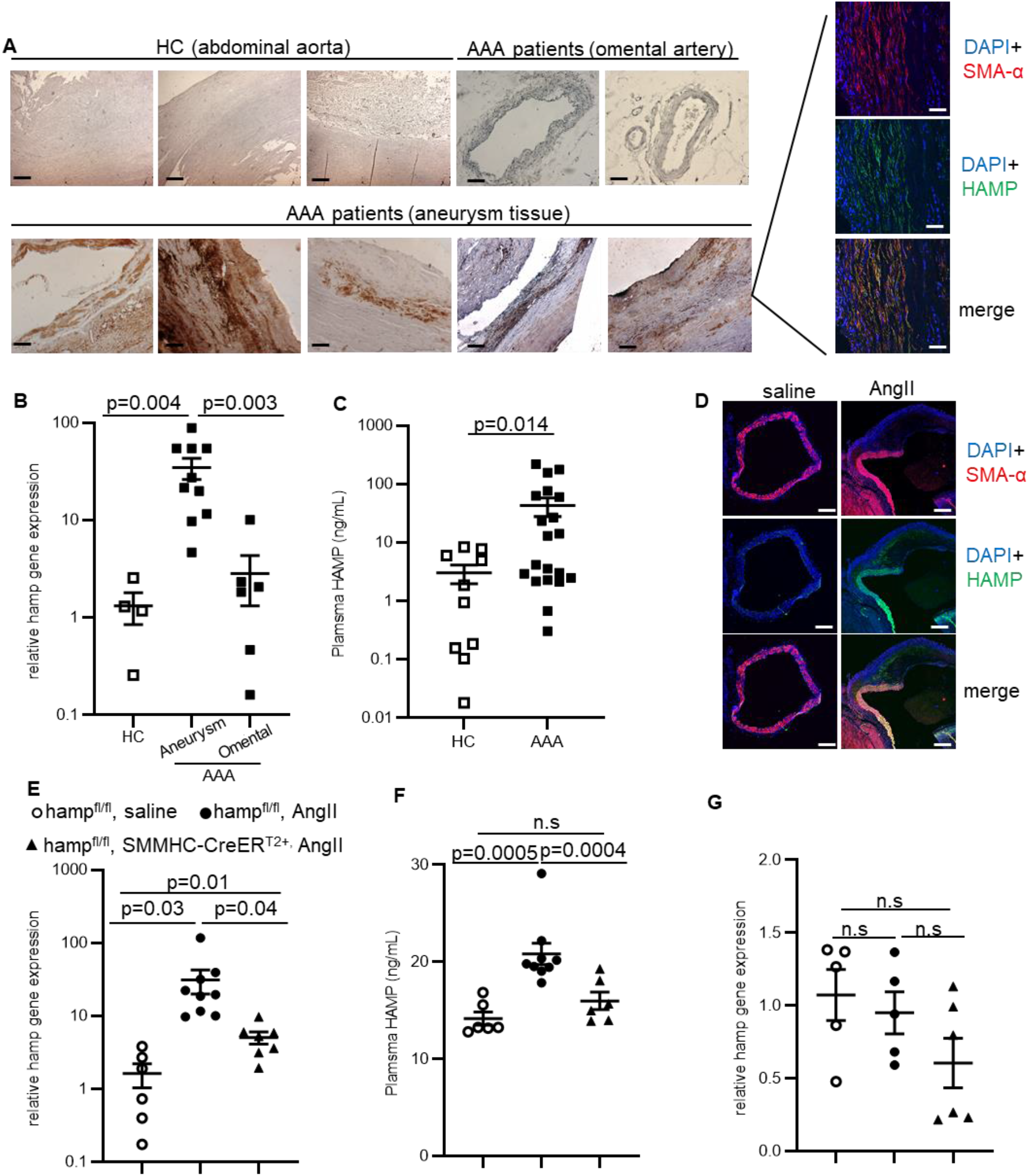
HAMP is elevated in SMCs of AAA in patients and in mice. **A)** HAMP immunostaining in abdominal aorta of healthy controls (HC), in omental artery and in aneurysm of abdominal aortic aneurysm (AAA) patients. **B)** Hamp gene expression in corresponding tissues. **C)** Plasma HAMP levels in HC and AAA patients. **D)** Repre-sentative HAMP immunostaining in aneurysm tissue of Ang-II-treated and control (sa-line) mice. **E)** Hamp expression in abdominal aorta of hamp^fl/fl^, saline-treated mice, hamp^fl/fl^, AngII-treated mice and hamp^fl/fl^, SMMHC-CreER^T2+^, AngII-treated mice. **F)** Plasma HAMP levels in corresponding animals. **G)** Hamp gene expression in livers of corresponding animals. Scale bar=100µM, original magnification x10. SMCs are iden-tified by staining for smooth muscle actin SMA-α. Values are shown at mean±S.E.M.

### SMC-derived HAMP is protective in the setting of AAA

Next, we set out to determine the role of SMC-derived HAMP in the pathophysiology of AAA. To that effect, we compared the effects of Ang-II treatment between Hamp^fl/fl^, SMMHC-CreER^T2+^ mice and Hamp ^fl/fl^ control mice. After 1 week, greater mortality with confirmed aneurysm rupture was observed in Hamp^fl/fl^, SMMHC-CreER^T2+^ mice than in Hamp ^fl/fl^ controls (Figure 2A). Amongst mice that survived to 1 week of Ang-II treatment, there was a greater incidence of non-fatal dissection and larger abdominal aortic lumen size in Hamp^fl/fl^, SMMHC-CreER^T2+^ mice than Hamp^fl/fl^ controls (Figure 2B, C). In AAA patients, higher plasma HAMP concentration at baseline was associated with lower aneurysm growth over the subsequent 12 months (Figure 2D). Together, these data demonstrate that raised HAMP in SMCs of the aneurysm tissue has a protective effect in the setting of AAA.

**Figure 2.**
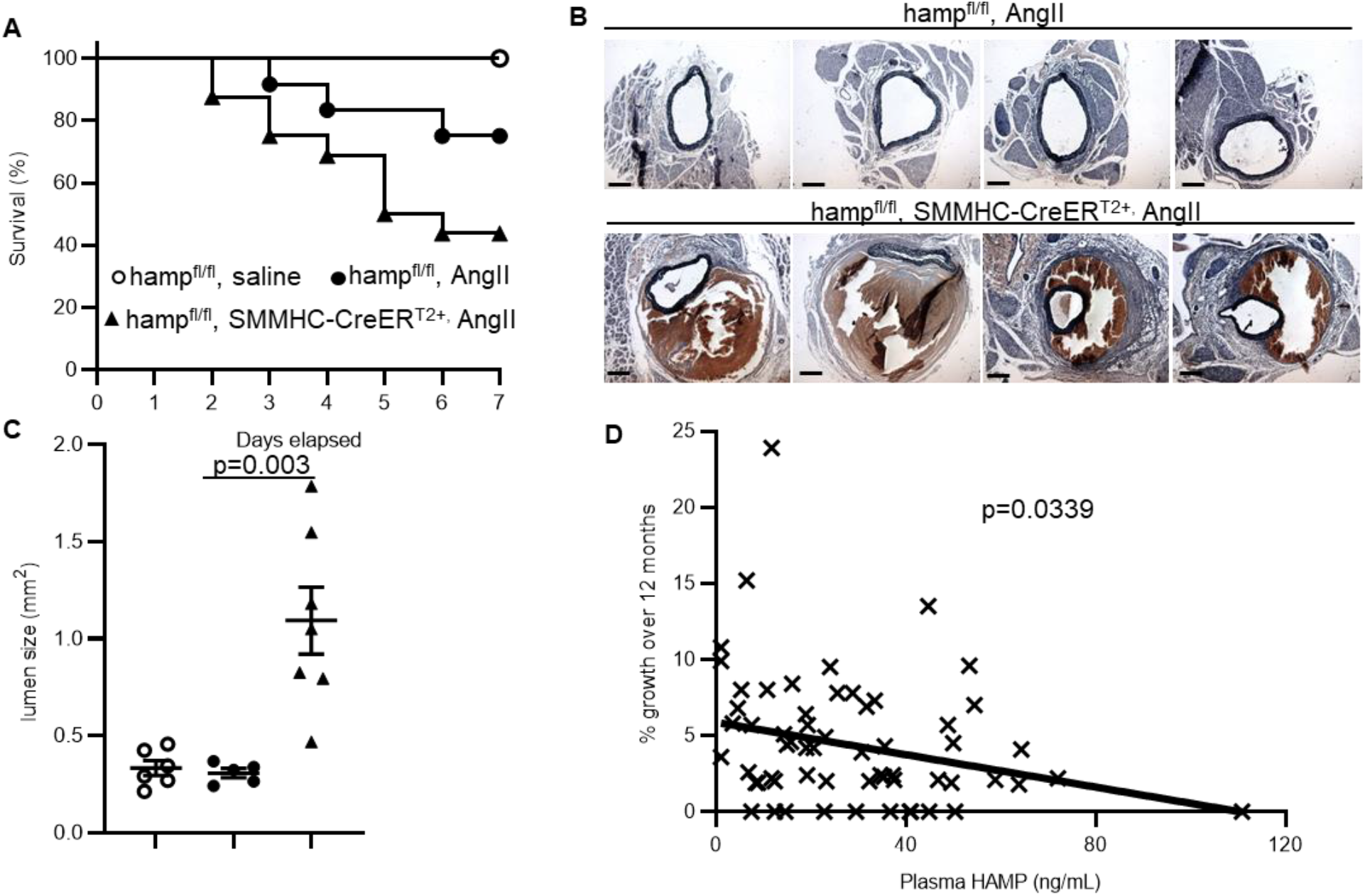
SMC-derived HAMP is protective in the setting of AAA. **A)** Seven-day survival of hamp^fl/fl^, saline-treated mice (n=6), hamp^fl/fl,^ AngII-treated mice (n=12) and hamp^fl/fl^, SMMHC-CreER^T2+^, AngII-treated mice (n=16). **B)** Representative images of elastin staining in abdominal aortas of hamp^fl/fl^, AngII-treated mice and hamp^fl/fl^, SMMHC-CreER^T2+^, AngII-treated mice. **C)** Lumen size of abdominal aortas from hamp^fl/fl^, saline-treated mice, hamp^fl/fl,^ AngII-treated mice and hamp^fl/fl^, SMMHC-CreER^T2+^, AngII-treated mice. **D)** Correlation between plasma HAMP levels at base-line and percentage (%) aneurysm growth over 12 months in AAA patients (n=62). Scale bar=200µM, original magnification x5. Values are shown at mean±S.E.M.

### The protective effect of SMC-derived HAMP is mediated by its cell-autonomous rather than its endocrine action

Having established that SMC-derived HAMP is protective in the setting of AAA, we next set out to determine whether such protection is mediated by its cell-autonomous or by its endocrine action. To explore the role of the cell-autonomous action of SMC-derived HAMP, we examined mice harbouring a conditional SMC-specific knock-in of HAMP-resistant ferroportin FPNC326Y (fpnC326Y^fl/fl^, SMMHC-CreER^T2+^). Ang-II treatment resulted in greater mortality with confirmed rupture in fpnC326Y^fl/fl^, SMMHC-CreER^T2+^ mice than in fpnC326Y^fl/fl^ controls (Figure 3A). Amongst the mice that survived to 1 week of Ang-II treatment, there was a greater incidence of non-fatal dissection and larger abdominal aortic lumen size in fpnC326Y^fl/fl^, SMMHC-CreER^T2+^ mice than in fpnC326Y^fl/fl^ controls (Figure 3B,C). Thus, loss of HAMP responsiveness in SMCs confers the same susceptibility to Ang-II-induced AAA as loss of SMC-derived HAMP itself. To explore the role of the endocrine action of SMC-derived HAMP, we examined markers of systemic iron homeostasis. We found that AngII treatment raised plasma iron levels (Figure 3D), reduced liver iron content (Figure 3E), and liver expression of the iron-regulated genes transferrin receptor1 Tfr1 and divalent metal transporter DMT1 (Figure 3F, G). These effects were maintained in Ang-II-treated fpnC326Y^fl/fl^, SMMHC-CreER^T2+^ mice, but not in Ang-II-treated hamp^fl/fl^, SMMHC-CreER^T2+^ mice (Figure 3D-G). These findings demonstrate that the endocrine action of SMC-derived HAMP contributes to Ang-II-driven changes in systemic iron indices, but in itself does not mediate the protective effect of HAMP in the setting of AAA. Thus, the protective effect of SMC-derived HAMP in the setting of AAA is dependent on its cell-autonomous action on FPN.

**Figure 3.**
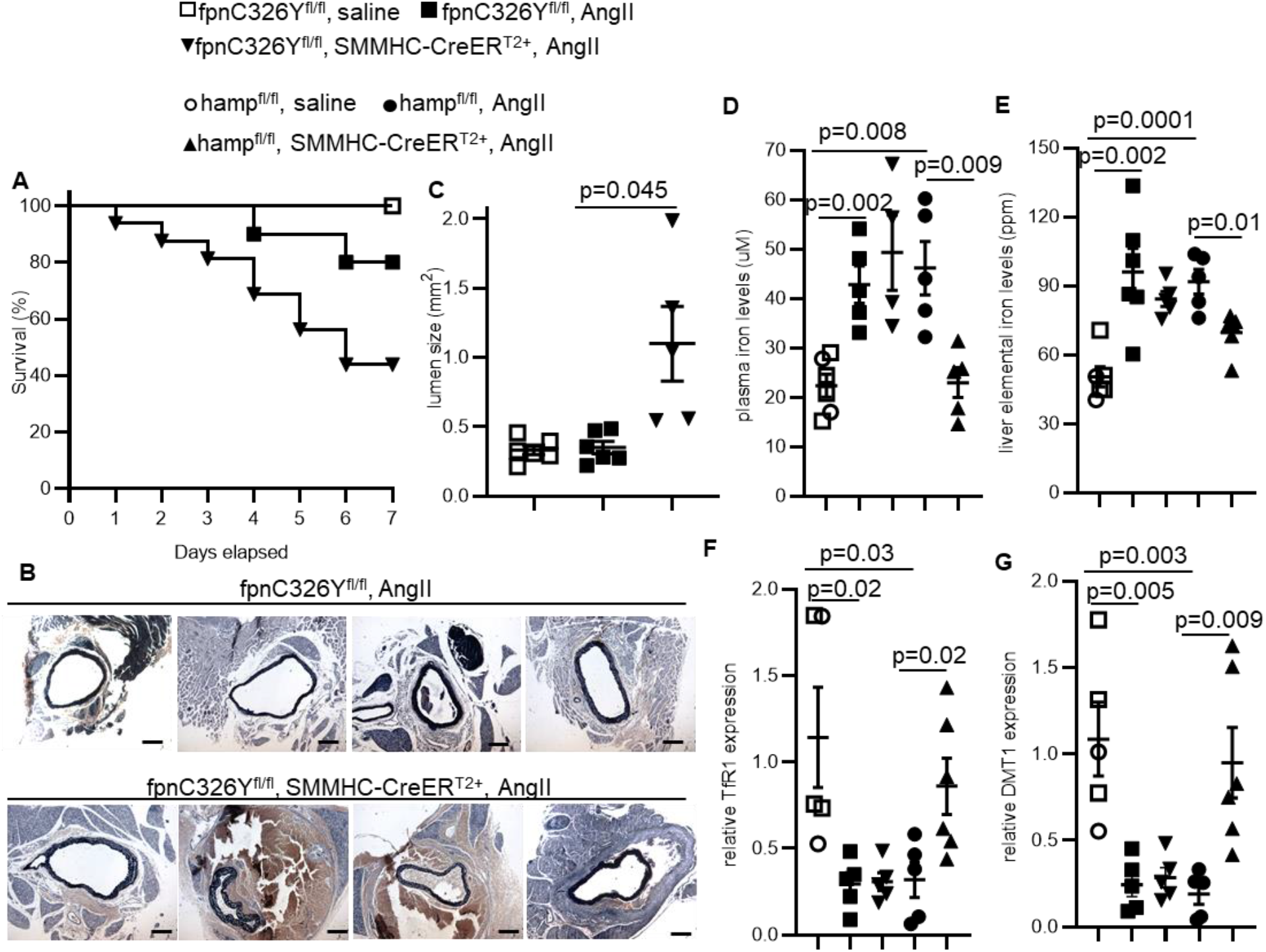
The protective effect of SMC-derived HAMP is mediated by its cell-autonomous rather than its endocrine action. **A)** Seven-day survival of fpnC326Y^fl/fl^, saline-treated mice (n=6), fpnC326Y^fl/fl^, AngII-treated mice (n=10) and fpnC326Y^fl/fl^, SMMHC-CreER^T2+^, AngII-treated mice (n=14). **B)** Representative images of elastin staining in abdominal aortas of fpnC326Y^fl/fl^, AngII-treated mice and fpnC326Y^fl/fl^, SMMHC-CreER^T2+^, AngII-treated mice. **C)** Lumen size of abdominal aortas from fpnC326Y^fl/fl^, saline-treated mice, fpnC326Y^fl/fl^, AngII-treated mice and fpnC326Y^fl/fl^, SMMHC-CreER^T2+^, AngII-treated mice. **D)** Plasma iron levels in fpnC326Y^fl/fl^, saline-treated mice, fpnC326Y^fl/fl^, AngII-treated mice, fpnC326Y^fl/fl^, SMMHC-CreER^T2+^, AngII-treated mice, hamp^fl/fl^, saline-treated mice, hamp^fl/fl^, AngII-treated mice and hamp^fl/fl^, SMMHC-CreER^T2+^, AngII-treated mice. **E)** Elemental iron concentrations in livers of corresponding mice (ppm=parts per million). **F)** Transferrin receptor TfR1 gene expression in livers of corresponding mice. **G)** Divalent metal transport DMT1 gene expression in livers of corresponding mice. Scale bar=200µM, original magnification x5. Values are shown at mean±S.E.M.

### Suppression of Lipocalin-2 mediates the protective effects of SMC-derived HAMP in the setting of AAA

Lipocalin-2 (LCN2) is an iron-scavenging protein known for its role in innate host defence^21^. LCN2 is expressed in many cell types, including smooth muscle cells, and involved in other processes, including promoting endoplasmic reticulum (ER) stress, cell death and suppressing autophagy^22-24^. We found that LCN2 expression is suppressed in aneurysms from AAA patients compared to abdominal aortic tissue from non-AAA healthy controls (HC), and to control vascular tissue (omental artery) from AAA patients (Figure 4A). In mice treated with AngII, LCN2 expression in the abdominal aorta was also suppressed in Hamp ^fl/fl^ mice. However, it was raised in hamp^fl/fl^, SMMHC-CreER^T2+^ mice (Figure 4B). Thus, suppression of local LCN2 expression in this setting requires SMC-derived HAMP. To determine whether failure to suppress LCN2 accounts for greater susceptibility of hamp^fl/fl^, SMMHC-CreER^T2+^ mice to AAA, we treated them with a LCN2-neutralising antibody at the start of Ang-II treatment. We found that LCN2-neutralising antibody reduced the incidence of fatal and non-fatal dissection and lowered abdominal aortic lumen size in these mice (Figure 4C, D, E). Thus suppression of local LCN2 expression mediates the protective effect of raised SMC-derived HAMP in the setting of AAA.

**Figure 4.**
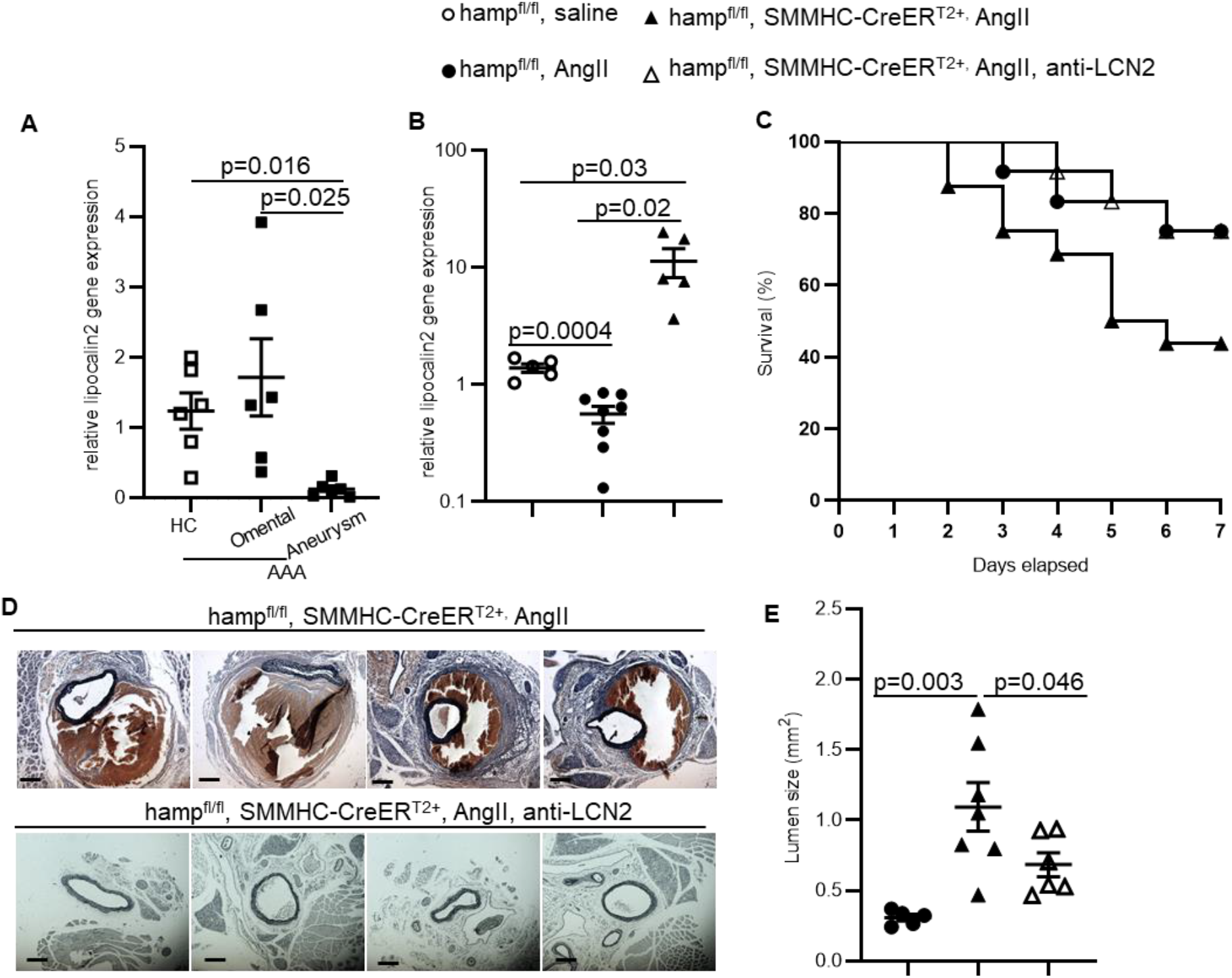
Suppression of LCN2 mediates the protective effects of SMC-derived HAMP in the setting of AAA. **A)** Lipocalin2 gene expression in abdominal aorta of healthy controls (HC), in omental artery and in aneurysm of abdominal aortic aneurysm (AAA) patients. **B)** Lipocalin2 gene expression in abdominal aortas of hamp^fl/fl^, saline-treated mice, hamp^fl/fl^, AngII-treated mice and hamp^fl/fl^, SMMHC-CreER^T2+^, AngII-treated mice. **C)** Seven-day survival of hamp^fl/fl^, AngII-treated mice (n=12), hamp^fl/fl^, SMMHC-CreER^T2+^, AngII-treated mice (n=16) and SMMHC-CreER^T2+^, AngII and LCN2-neutralising antibody (anti-LCN2)-treated mice (n=12). **D)** Representative images of elastin staining in abdominal aortas of corresponding mice. **E)** Lumen size of abdominal aortas in corresponding mice. Scale bar=200µM, original magnification x5. Values are shown at mean±S.E.M.

## Discussion

The most important finding of the present study is that the cell-autonomous action of SMC-derived HAMP is protective in the setting of AAA. Indeed, loss of HAMP specifically in SMCs increased the incidence of fatal and non-fatal dissection in an experimental mouse model of Ang-II-induced AAA. This phenotype was replicated in mice with loss of HAMP responsiveness specifically in SMCs. Consistent with a protective role for HAMP, higher plasma HAMP levels are associated with slower AAA growth in patients.

Another important finding is that SMC-derived HAMP in the aneurysm tissue contributes to raising plasma HAMP levels. Indeed, plasma HAMP levels were raised in an experimental model of AAA, but only in mice with intact HAMP expression in SMCs. These findings are the first formal demonstration of the contribution of ectopic (non-hepatic) HAMP to plasma HAMP levels.

The mechanisms underlying the increase in HAMP expression within the aneurysm tissue are not understood. However, the aneurysm tissue is an inflammatory microenvironment and it is well recognised that HAMP is induced by inflammation, specifically interleukin 6^25^. Additionally, the aneurysm tissue is characterised by iron deposition, likely deriving from the intraluminal thrombus^13^. Indeed, we found that AAA tissue from patients displayed heavy iron deposits particularly in the SMC layer (supplemental figure 1). When we treated vascular smooth muscle cells with excess iron in-vitro (in the form of ferric citrate) or with interleukin-6, we observed a rapid induction of *hamp* gene expression (supplemental figure 2), supporting the notion that both local iron deposition and inflammation may account for raising HAMP expression in SMCs within the aneurysm tissue.

The mechanisms underlying the protective effect of SMC-derived HAMP appear to depend, at least in part, on the suppression of local LCN2 expression. Indeed, LCN2 expression was reduced in patient AAA tissue. A similar effect was seen in mouse model of AAA but only in mice with intact HAMP expression in SMCs. LCN2 is best known for its role in innate immunity, mediated by sequestering iron away from bacteria^21^. It is expressed in neutrophils and at lower levels in the other cells including the renal tubules, cardiomyocytes, neurons and smooth muscle cells^22-24^. In cardiomyocytes and neurons, it has been shown that LCN2 mediates cell death by inhibiting reparative autophagy and promoting apoptosis^24^. Autophagy is also recognised as a protective mechanism that maintains tissue integrity in the setting of AAA^25,26^. Additionally, LCN2 has neutrophil chemotactic properties, and neutrophil infiltration is a recognised mechanism of tissue injury in the setting of AAA^27-29^. Thus, HAMP-mediated suppression of LCN2 could act to dampen cell death and reduce neutrophil infiltration in AAA (Figure 5).

**Figure 5.**
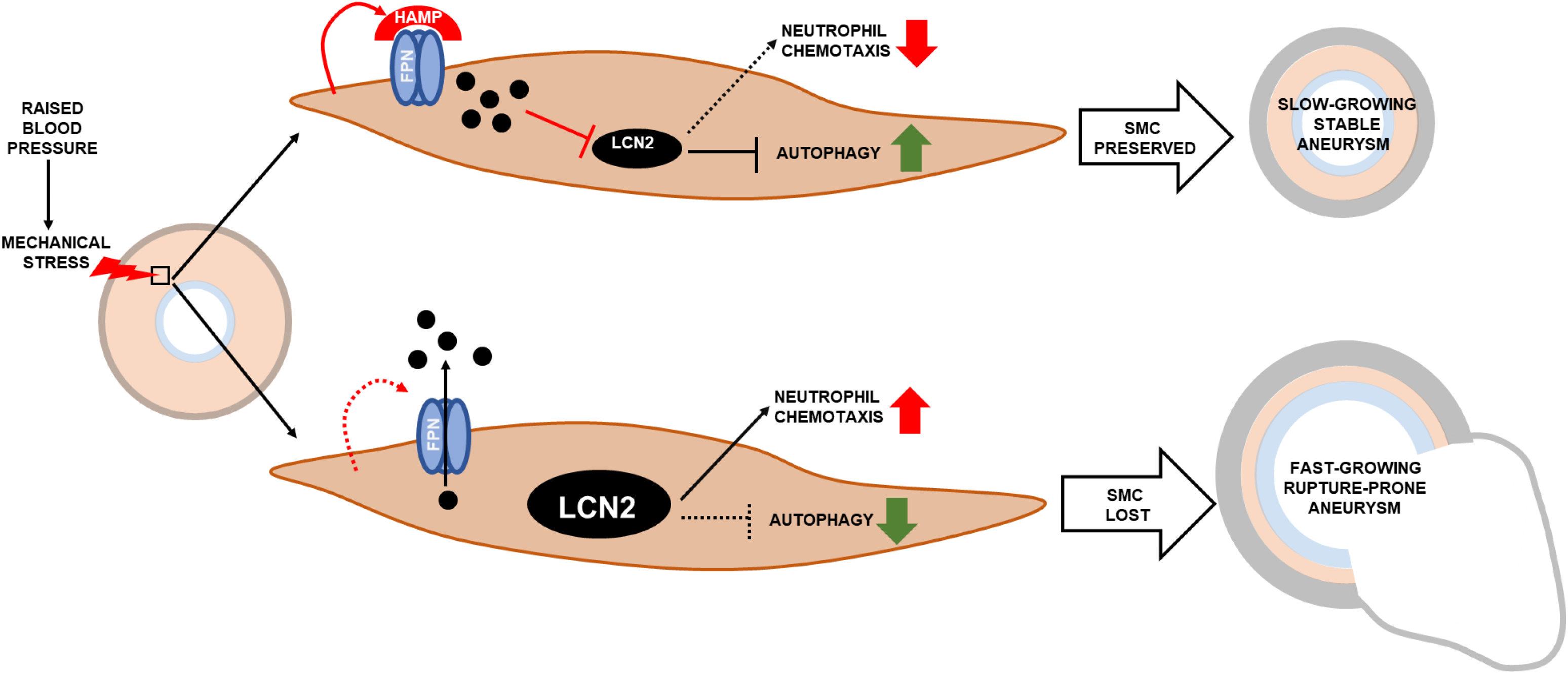
Model for the Protective Role of Smooth Muscle Cell HAMP in AAA. During the development of AAA, expression of HAMP (HAMP) in smooth muscle cells (SMCs) of the abdominal aorta is raised, causing cell-autonomous inhibition of ferroportin (FPN) and increasing intracellular iron retention. Lipocalin 2 (LCN2) normally promotes AAA growth by inhibiting autophagy in SMCs and attracting neutrophils. HAMP-mediated intracellular iron retention inhibits LCN2 expression, and in doing so helps preserve SMCs in the tunica media. Preservation of SMCs results in slower growing/more stable aneurysms.

Since the discovery of HAMP at the turn of the century, the consensus has been that it derives primarily from the liver and operates solely as an endocrine regulator of systemic iron homeostasis. In recent years, work from this team has offered new understanding by demonstrating that ectopic HAMP found in other tissues operates in an autocrine fashion to control local iron homeostasis in a manner that is important for the normal physiological function of those tissues (6, 7, 10). The present study adds a new dimension to that understanding, by demonstrating a disease-modifying role for ectopic HAMP. In doing so, it highlights not only the multi-faceted roles of HAMP in normal and pathophysiology, but also its potential as a therapeutic target in conditions other than disorders of iron homeostasis.

## ACKNOWLEDGEMENT

We acknowledge the support from the Oxford Transplant Biobank (Sandrine Rendel, Jon Milton and Prof Rutger Ploeg) to enable access to non-aneurysm aortic tissue samples collected for the biobank.

## FIGURE LEGENDS

**Supplemental Figure 1.**
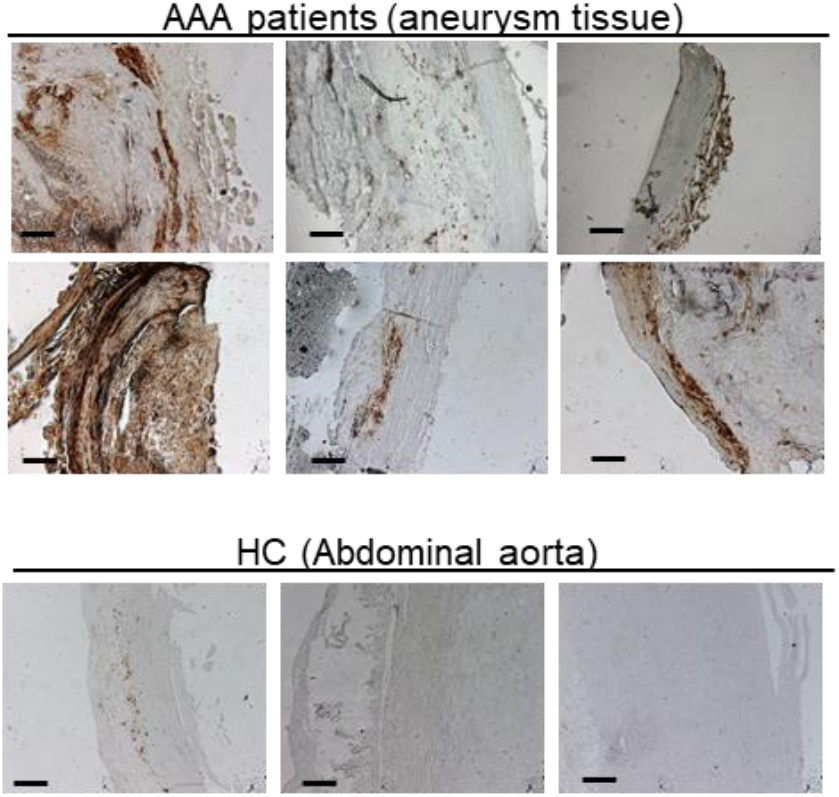
AAA is characterised by iron deposition in smooth muscle cells of the abdominal aorta. Representative images of DAB-enhanced Peris iron stain in abdominal aorta of healthy controls (HC), in omental artery and in aneurysm of abdominal aortic aneurysm (AAA) patients. Scale bar=200μM, original magnification x5.

**Supplemental Figure 2.**
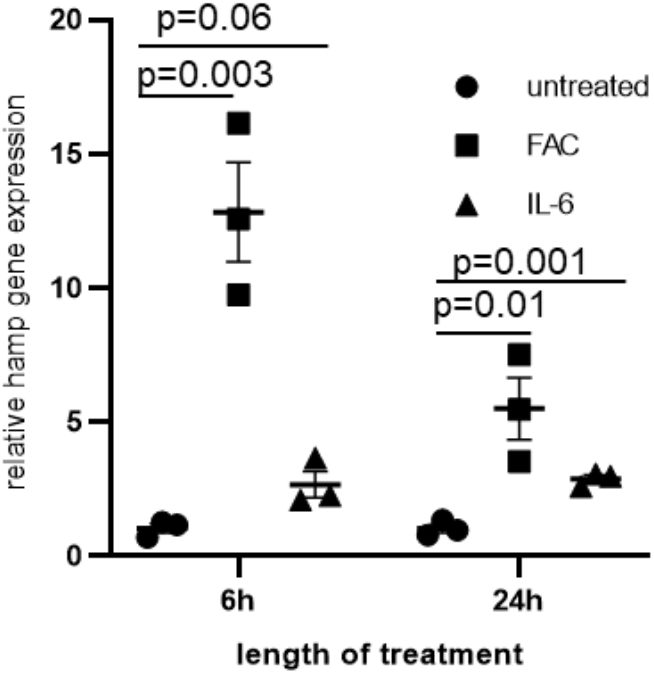
Hepcidin gene expression is induced by iron and interleukin-6 in vascular smooth muscle cells. Mouse vascular smooth muscle cells were treated with 500μM ferric citrate (FAC) or 100ng/mL recombinant human interleukin-6 (IL-6) for 6 or 24 hours. Values are shown at mean±S.E.M.

